# A Scalable Topical Vectored Vaccine Candidate Against SARS-CoV-2

**DOI:** 10.1101/2020.05.31.126524

**Authors:** Mohammed A Rohaim, Muhammad Munir

**Affiliations:** Division of Biomedical and Life Sciences, Faculty of Health and Medicine, Lancaster University, Lancaster LA1 4YG, UK

## Abstract

The severe acute respiratory syndrome-coronavirus 2 (SARS-CoV-2) caused an ongoing unprecedented global public health crises of coronavirus disease in 2019 (CoVID-19). The precipitously increased death rates, its impact on livelihood and trembling economies warrant the urgent development of SARS-CoV-2 vaccine which would be safe, efficacious and scalable. Owing to unavailability of the vaccine, we propose a *de novo* synthesised avian orthoavulavirus 1 (AOaV-1)-based topical respiratory vaccine candidate against CoVID-19. Avirulent strain of Newcastle disease virus, proto-type virus of AOaV-1, was engineered to express full length spike (S) glycoprotein which is highly neutralizing and major protective antigen of the SARS-CoV-2. Broad-scale *in vitro* characterization of recombinant vaccine candidate demonstrated efficient co-expression of the hemagglutinin-neuraminidase (HN) of AOaV-1 and S protein of SARS-CoV-2, and comparable replication kinetics were observed in cell culture model. The recombinant vaccine candidate virus actively replicated and spread within cells independently of exogenous trypsin. Interestingly, incorporation of S protein of SARS-CoV-2 into the recombinant AOaV-1 particles attributed the sensitivity to anti-SARS-CoV-2 antiserum and more prominently to anti-AOaV-1 antiserum. Finally, our results demonstrated that the recombinant vaccine vector stably expressed S protein after multiple propagation in chicken embryonated eggs, and this expression did not significantly impact the *in vitro* growth characteristics of the recombinant. Taken together, the presented respiratory vaccine candidate is highly attenuated in primates *per se*, safe and lacking pre-existing immunity in human, and carries the potential for accelerated vaccine development against CoVID-19 for clinical studies.

## Introduction

An outbreak of pneumonia was erupted in Chinese seafood market in Wuhan during late 2019 and within a month of its origin, on 30 January 2020, a Public Health Emergency of International Concern was declared by the World Health Organization (WHO) due to its high human-to-human transmission. Within next month, the outbreak of coronavirus disease in 2019 (CoVID-19) soared among communities unprecedentedly and spread across the globe and a pandemic was declared on 11 March 2020 by WHO. A large proportion (∼80%) of CoVID-19 infected patients showed only moderate symptoms which led to staggering rate increase in global spread of the infection. The acute respiratory distress syndrome, manifested in ∼20% of CoVID-19 patients, caused substantial case fatality rates especially in elderly and frail people with co-morbidities^1^.

The severe acute respiratory syndrome-coronavirus 2 (SARS-CoV-2), the causative agent of ongoing CoVID-19 pandemic, belongs to the family *Coronavirdae* within the *Betacoronavirus* (β-CoV) genus. Similar to SARS-CoV and Middle Eastern respiratory syndrome-related coronavirus (MERS-CoV), SARS-CoV-2 carries a single stranded linear RNA genome with positive polarity^2^. The genome is consisted of 4 structural proteins including envelope (E), spike (S), membrane (M), and nucleocapsid (N), 16 non-structural proteins (nsp 1-16) and multiple accessary proteins. Amongst these viral proteins, the S protein constitute a major protective antigen that elicit highly specific antibodies mediated immune responses^2^. Therefore, the S protein remained the primary vaccine markers against coronaviruses.

Currently, there is no registered drugs or vaccines available to curb the pandemic; however, multiple vaccines using a range of technologies are being developed, or pre-clinically or clinically being investigated^3^. Amongst these, an inactivated vaccine has elicited strong antibodies which can neutralize multiple SARS-CoV-2 strains and can partially or fully protect macaques against SARS-CoV-2 challenge^4^. A chimpanzee adeno (ChAd)-vectored vaccine, expressing the full-length S gene of SARS-CoV-2, elicited humoral and cell-mediated responses in rhesus macaques^5^. However, it failed to fully alleviated clinical signs in vaccinated macaques albeit reduced severity and protection against pneumonia. The ChAdOx1 nCoV-19 also failed to reduce the viral replication in the nose, highlighting the potential spread of SARS-CoV-2 through sneezing even in vaccinated people^5^.

Owing to multifaceted advantages, the avian orthoavulavirus 1 (AOaV-1) proposes a potential vaccine vector against SARS-CoV-2. Specifically, AOaV-1 (represented by a type species, Newcastle disease virus, NDV) are exclusively cytoplasmic viruses and therefore the viral gene segments are not integrated into the host genome which raises their safety profile. Since these vectors lack natural recombination, the expression of transgenes is genetically stable. Additionally, AOaV-1 can infect multiple species of animals, the vaccines can be produced in chicken embryonated eggs and multiple cell lines^6^. Given these and other features, apathogenic strains of AOaV-1 have been used as live attenuated vaccines against multiple viruses including influenza, SARS, human immunodeficiency virus^7,8^, human parainfluenza, rabies^9^, Nipah disease^10^, Rift Valley fever^11^, Ebola and highly pathogenic H5N1^12,13^. Importantly, AOaV-1 has appeared to be safe and effective in mice ^10,13^, dogs^9^, pigs^10^, cattle^14,15^, sheep^14^, African green and rhesus monkeys^16,17^ and humans ^18-21^. Notably, the natural host of AOaV-1 are birds and the vector is antigenically distinct from common human pathogens. Therefore, no pre-existing immunity in human make it an ideal vector to deliver transgene effective and safely.

In the present study, we *de novo* designed an AOaV-1 vector and generated a recombinant vaccine candidate by expressing full-length codon optimized S protein of the SARS-CoV-2 at a pre-optimized gene junction. This topical respiratory vaccine candidate was fully characterized *in vitro*. Based on the infection and spreadability within cells, its sensitivity to anti-AOaV-1 and anti-SARS-CoV-2 neutralizing antibodies, its replication kinetics and stability in chicken embryonated eggs, the rAOaV-1-SARS-CoV-2 is a scalable topical vectored vaccine candidate against SARS-CoV-2 to be tested for safety and immunogenicity in animal studies.

## Results

### Design and Construction of AOaV-1-SARS-CoV-2 Vaccine Candidate

An avirulent strain of AOaV-1 was used to construct vaccine candidate against SARS-CoV-2. The full length antigenomic sequence of AOaV-1, originally isolated from asymptomatic wild birds, was *de novo* synthesised. To facilitate minigenome transcription, an autocatalytic and “rule-of-six” adhered hammerhead ribozyme sequences was introduced in both 5’ and 3’-ends of the antigenome (**Fig. 1A**).

**Figure 1.**
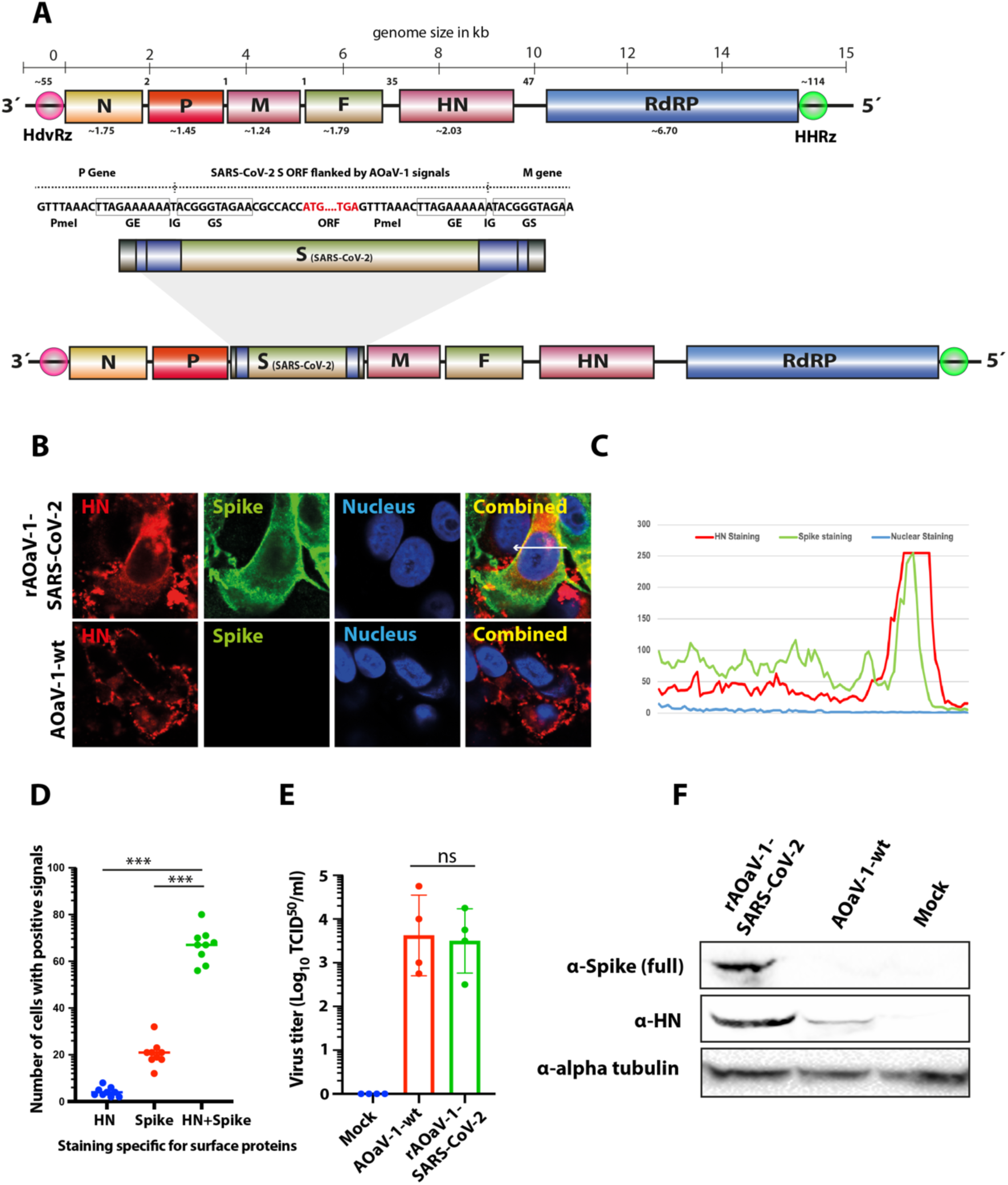
Construction, rescue and characterization of rAOaV-1-SARS-CoV-2 vaccine candidate. **(A)** The full-length ORF for S gene of SARS-CoV-2 was over-hanged with required transcriptional signals (GE, GS, IG) and inserted in between P and M genes. The rough gene size is mentioned below each gene, the division of the genome across the length and number of nucleotides in intergenic region is displayed at the top of the schema of the AOaV-1 genome. (**B**) Vero cells were infected with the AOaV-1-wt or rAOaV-1-SARS-CoV-2 and sainted for the expression of the HN (red) or S (green) proteins. The co-expression of both surface proteins is coloured yellow in combined images. (**C**) Quantitative co-expression profile is marked with arrow and shown in the line chart. (**D**) A total of 9 microscopic fields were scanned for the presence of HN or S or both proteins. A significantly higher proportion of HN+S expressing cells was identified. (**E**) A comparable replication of AOaV-1-wt or rAOaV-1-SARS-CoV-2 in Vero cells indicating the competitive replication of AOaV-1 even after the expression of the foreign S gene. (**F**) Western blot analysis for the expression of the HN protein indicating the active replication of the AOaV-1 and S protein indicating the replication competence of the recombinant virus, rAOaV-1-SARS-CoV-2. Alpha tubulin was used as loading control. The asterisk indicates the level of significance, whereas ns indicate non-significance.

An expression cassette containing the Kozak sequence, GE, GS, IG and the ORF for the full-length S gene was codon-optimized for *homo sapiens* codon usage and inserted into the unique *PmeI* site between P and M genes, which was originally preserved during the cloning of complete antigenome (**Fig. 1A**). The construct was named rAOaV-1-SARS-CoV-2 whereas the AOaV-1 without the insertion of the foreign gene (AoaV-1-wt) was used as infection control throughout the study. The orientation of the inserted S gene was confirmed by the nucleotide sequence analysis.

### Rescue of Recombinant Vaccine and Evaluation of the Spike Gene Expression of SARS-CoV-2

The rAOaV-1-SARS-CoV-2 and AOaV-1-wt were rescued in Vero cells and propagated in 8-day-old embryonated chicken eggs. Screening of multiple individually inoculated eggs, using real-time PCR and hemagglutination assays, has identified successfully rescued rAOaV-1-SARS-CoV-2 viruses (**Fig. 1 Supplementary**) which were used to fully characterize in the presented study.

Expression of the S protein was confirmed by indirect confocal immunofluorescent staining of rAOaV-1-SARS-CoV-2-infected Vero cells. Counter-staining of the HN protein of AOaV-1 and S protein of SARS-CoV-2 confirmed co-expression of the surface protein in rAOaV-1-SARS-CoV-2-infected Vero cells, whereas only HN protein expression was observed in AOaV-1-wt-infected cells (**Fig. 1B**). Graphical profile of the expression intensities confirmed co-expression of both surface proteins (**Fig. 1C**) where a vast majority of rAOaV-1-SARS-CoV-2-infected cells expressed both HN and S proteins simultaneously (**Fig. 1D**).

Both recombinant and wild type AOaV-1 isolates replicated at high titre in eggs (≥28 HAU/ml, data not shown). The evaluation of the viral replication in the presence of exogenous proteases in Vero cells indicated that rAOaV-1 expressing codon optimized S gene replicated at the level comparable to wild type AOaV-1 (**Fig. 1E**). The expression analysis of the S protein by Western blot indicated a potent expression of the full-length S protein in rAOaV-1-SARS-CoV-2-infected cells whereas expression of the HN proteins was detected in both recombinant and wt AOaV-1-infected cells (**Fig. 1F and Fig. 3 supplementary**). These results confirm that the expression of the transgene (S) didn’t interfere with the growth characteristics of the AOaV-1 and could be a replication competitive vaccine candidate.

### Exogenous Trypsin Independent Growth Characteristics of Vaccine Construct

To initiate virus replication, the F protein of the AOaV-1 has to be cleaved by cellular proteases^22^. In order to investigate the pre-requisite of exogenous trypsin-like extracellular proteases for the infectivity of AOaV-1, eggs-propagated rAOaV-1-SARS-CoV-2 and AOaV-1-wt were used to infect Vero cells with a multiplicity of infection (MOI) of 1.0 without exogenous trypsin treatment. As expected, the replication of rAOaV-1-SARS-CoV-2 was apparent 6 hours post-infection and the individually infected cells spread the infection to neighbouring cells within next 12 hours. At 2 days post-infection, as high as 90% of the cells were infected with the rAOaV-1-SARS-CoV-2 and majority of cells co-expressed HN protein of the AOaV-1 and S protein of the SARS-CoV-2 (**Fig. 2A and Fig. 2 supplementary**). Cumulative fluorescence dynamics, based on either HN or S protein expressions, over the course of the infection, confirmed a gradual spread of the infection (**Fig. 2B**). Co-expression fluorescence profile highlighted the co-localization of both surface proteins (**Fig. 2C**). Similar to recombinant AOaV-1, the AOaV-1-wt replicated to a similar extent over the course of two days post-infection without the need of exogenous extracellular proteases and saturated level of expression of the HN protein was observed at 48 hours post-infection (**Fig. 2D and Fig. 2 supplementary**). The cumulative fluorescence dynamics (**Fig. 2E**) and fluorescence profile (**Fig. 2F**) confirmed the active and progressive expression and HN-specific staining, respectively. Taken together, these results confirm the active replication, and expression of AOaV-1 and foreign genes in Vero cells independently of exogenous trypsin.

**Figure 2.**
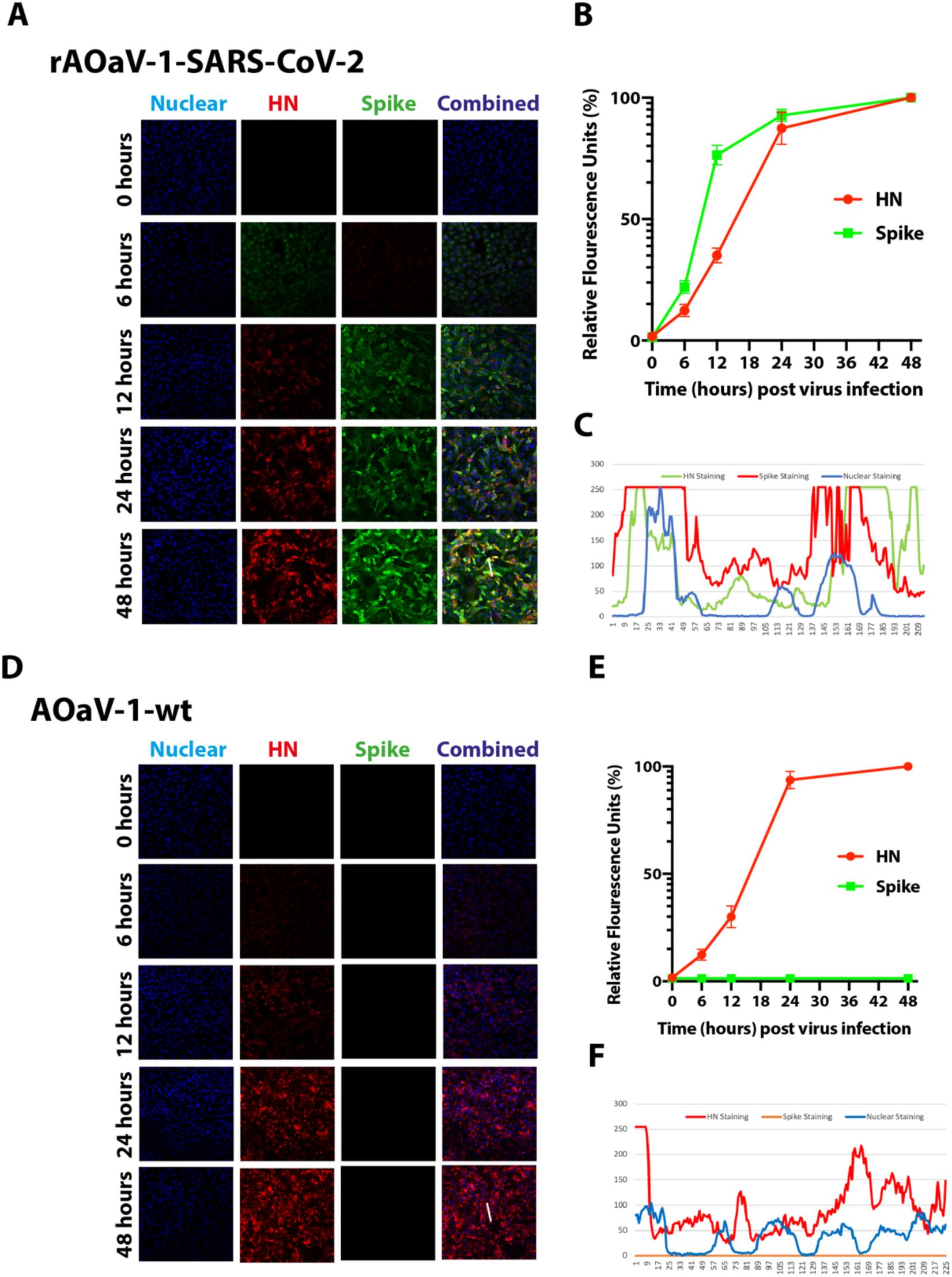
Replication of rAOaV-1-SARS-CoV-2 and AOaV-1-wt in mammalian cells. **(A)** Vero cells were infected with moi of 1.0 with rAOaV-1-SARS-CoV-2. These cells were fixed with paraformaldehyde at 0, 6, 12, 24- and 48-hours post-infection and stained for the HN protein of rAOaV-1 and S protein in the rAOaV-1-SARS-CoV-2. (**B**) Cumulative quantification of the green (S) and red (HN) fluorescence intensities before confocal microscopic imaging. (**C**) The co-expression profile for the HN and S proteins. **(D)** Vero cells were infected with moi of 1.0 with AOaV-1-wt. These cells were fixed with paraformaldehyde at 0, 6, 12, 24- and 48-hours post-infection and stained for the HN protein of AOaV-1. (**E**) Cumulative quantification of the green (S) and red (HN) fluorescence intensities before confocal microscopic imaging. (**F**) The graphically presentation of co-expression profile for the HN and S proteins.

**Figure 3.**
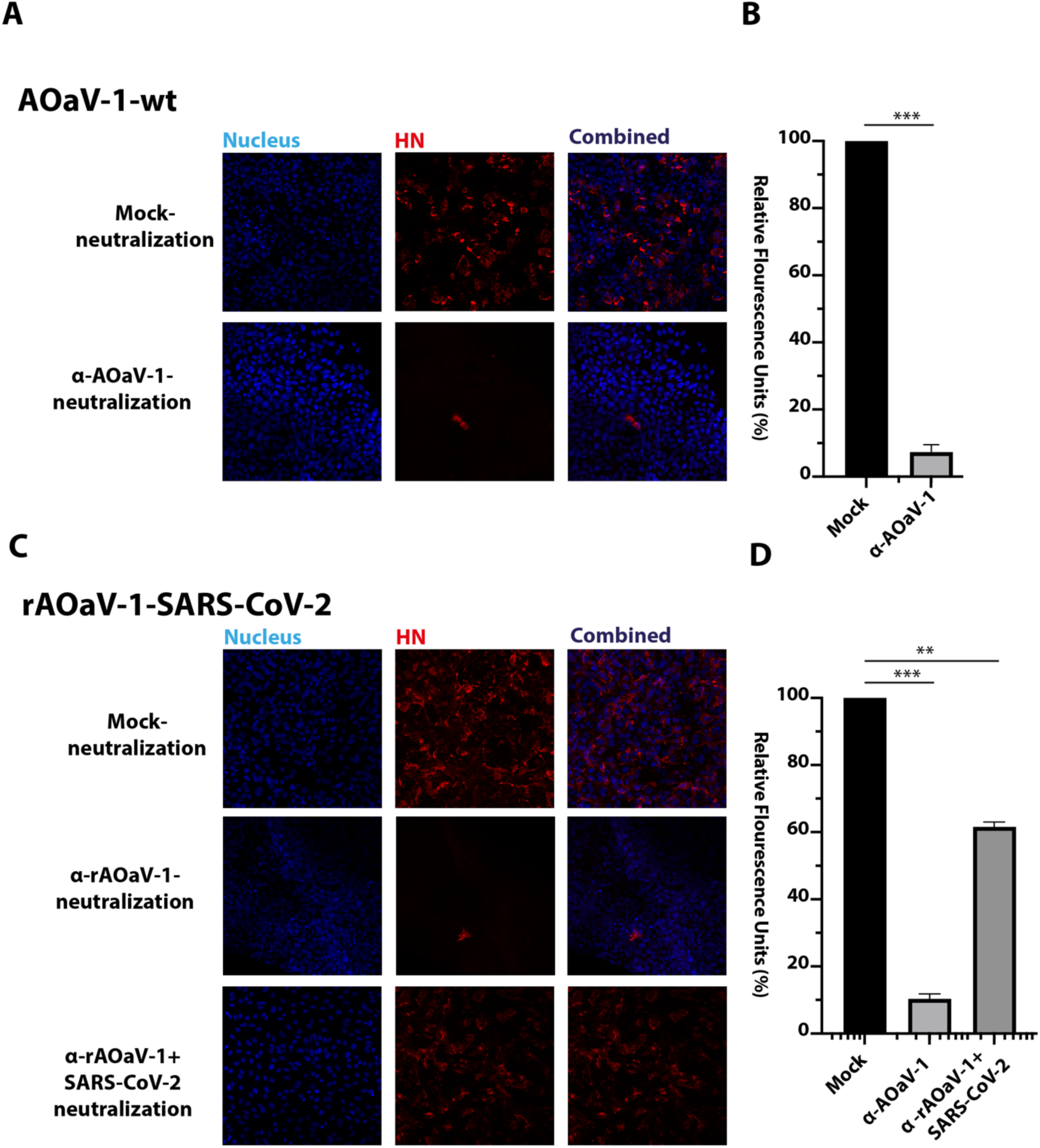
Sensitivity of recombinant and wt viruses against antisera. **(A)** AOaV-1-wt virus was incubated with an antiserum against AOaV-1 for 2 hours or was incubated with plan antiserum. The Vero cells were then infected with these viruses and incubated for 24 hours before staining against the HN protein of AOaV-1. (**B**) The mock-neutralization was set to 100%, and the quantitative analysis of the virus replication was plotted for the AOaV-1 neutralized with antiserum. (**C**) rAOaV-1-SARS-CoV-2 virus was incubated with an anti-AOaV-1 antiserum or SARS-CoV-2 antiserum for 2 hours before infection of Vero cells for another 24 hours. These cells were then stained for the expression of the HN glycoprotein of rAOaV-1-SARS-CoV-2. (**D**) Quantitative measurement of the staining intensities plotted against the mock-treated neutralization control. The asterisk indicates the level of significance.

### The rAOaV-1-SARS-CoV-2 is Sensitive to AOaV-1 and SARS-CoV-2 Antisera

The F and HN surface glycoproteins are critical for receptor binding as well as membrane fusion which are indispensable for virus entry and subsequent initiation of the virus replication^22,23^. In contrast, the coronaviruses have only a single enveloped glycoprotein which perform dual functions of receptor binding and membrane fusion^24^. In order to understand the influence of the S protein of the SARS-CoV-2 on the infectivity of recombinant virus, the sensitivities of AOaV-1 and SARS-CoV-2 antisera were assessed and compared for neutralization. As expected, the AOaV-1-wt was resistant to mock antiserum neutralization and was almost fully neutralized by the serum against the AOaV-1 (**Fig. 3A**). Quantitatively, the anti-AOaV-1 antiserum reduced the infection of AOaV-1-wt in Vero cells by ∼92% compared to the mock-neutralization (**Fig. 3B**). Neutralization analysis of the rAOaV-1-SARS-CoV-2 showed a significant blockage of the virus replication by pre-incubation and subsequent infection of rAOaV-1-SARS-CoV-2 with either anti-AOaV-1 antiserum or anti-SARS-CoV-2 anti-serum (**Fig. 3C**). There was no neutralization observed upon virus’s treatment with the naïve mouse control serum. Compared to mock-neutralization, as high as 90% inhibition of the rAOaV-1-SARS-CoV-2 was observed with anti-AOaV-1 antiserum and ∼40% inhibition was noticed using anti-SARS-CoV-2 anti-serum (**Fig. 3D**). These observations confirm that the incorporation of S protein of SARS-CoV-2 into the recombinant AOaV-1 particles attributed the sensitivity of the AOaV-1 to anti-SARS-CoV-2 antiserum and anti-AOaV-1 antiserum.

### In Vitro Growth Characterization of the Vaccine Construct

In order to understand the replication competence of rAOaV-1-SARS-CoV-2 and AOaV-1-wt, a multistep growth kinetics was evaluated in Vero cells. The time-course quantitative measurement of the genomic copies confirmed that the expression of the S gene didn’t interfere with the viral replication and the rAOaV-1-SARS-CoV-2 replicated competitively and comparability with the AOaV-1-wt (**Fig. 4A**). The cell lysate from same experimental setting was used to assess the expression of the AOaV-1 structural proteins. The expression analysis using Western blotting demonstrated that both AOaV-1-wt (**Fig. 4B**) and rAOaV-1-SARS-CoV-2 (**Fig. 4C**) progressively replicated in Vero cells and expressed the HN protein as early as 6 hours post-infection and as late as 2 days after initiation of the infection. These results demonstrated that the expression of the foreign genes by the AOaV-1 was unable to further attenuate the replication of the virus and could yield into sufficient viral quantity in cell culture system.

**Figure 4.**
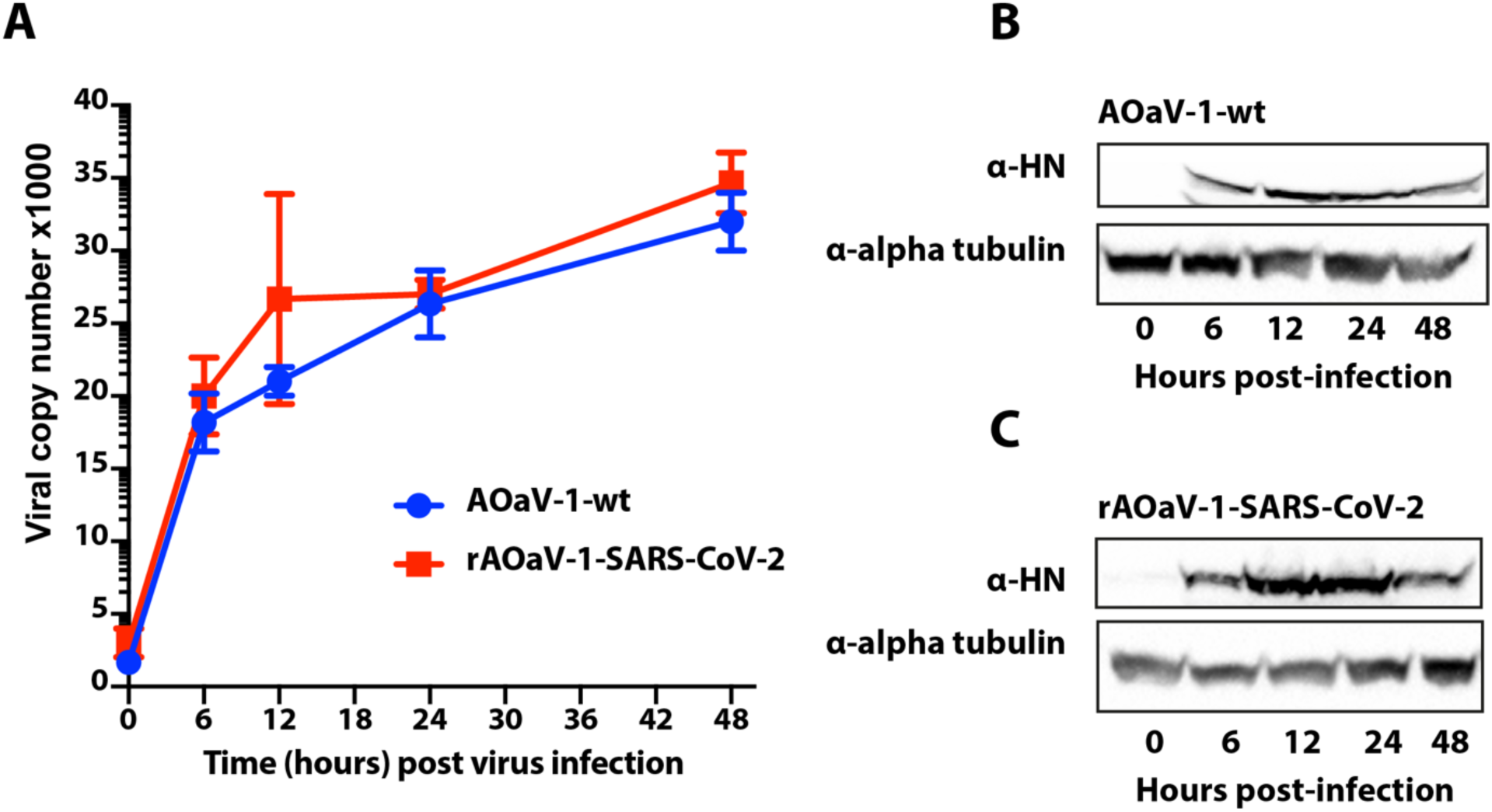
Growth kinetics of the vaccine and wt constructs. **(A)** Vero cells were infected with an moi 1.0 of rAOaV-1-SARS-CoV-2 or AOaV-1-wt for 2 hours (0 hour post infection) and were then replaced with media to be incubated and extraction of RNA at 6, 12, 24, 48 hours post infection. The viral copy number were calculated from AOaV-1 standard run in parallel. (**B**) Vero cells were infected with AOaV-1 and total cell lysate was prepared at indicated time points post-infection. These lysates were run for Western blotting using HN antibodies to demonstrate virus replication and alpha tubulin as loading control. (**C**) Similar to section B, cells were infected with rAOaV-1-SARS-CoV-2 and the expression for the HN and alpha tubulin was measured at indicated time points.

### Stability of Recombinant AOaV-1 SARS-CoV-2 Vaccine Candidate

In order to assess the stability of the foreign gene in the recombinant AOaV-1, the recovered and S-gene expression-confirmed viruses were passaged in 8-days-old embryonated chicken eggs for five consecutive passages. Both rAOaV-1-SARS-CoV-2 and AOaV-1-wt replicated substantially in eggs (≥28 HAU/ml, data not shown). The comparative replication competence was assessed between first and fifth eggs passaged viruses in Vero cells. The rAOaV-1-SARS-CoV-2 expressing the S protein of the SARS-CoV-2 grew efficiently and at the level of the AOaV-1-wt in the first passage as well as after 5^th^ passage in the embryonated eggs (**Fig. 5A**). Correspondingly, the expression of the structural protein of the AOaV-1 further confirmed the stable propagation of the wt and recombinant viruses at least for several passages (**Fig. 5B** and **Fig. 3 supplementary**). The sequence integrity of the inserted S gene as well as the P and M junction was assessed without any mutations. Additionally, sequencing of the S gene from the first and fifth passages confirmed no unwanted mutations in purified viruses. These results demonstrate that the rAOaV-1-SARS-CoV-2 expresses S gene stably and this expression did not significantly impact the *in vitro* growth characteristics of the recombinants.

**Figure 5.**
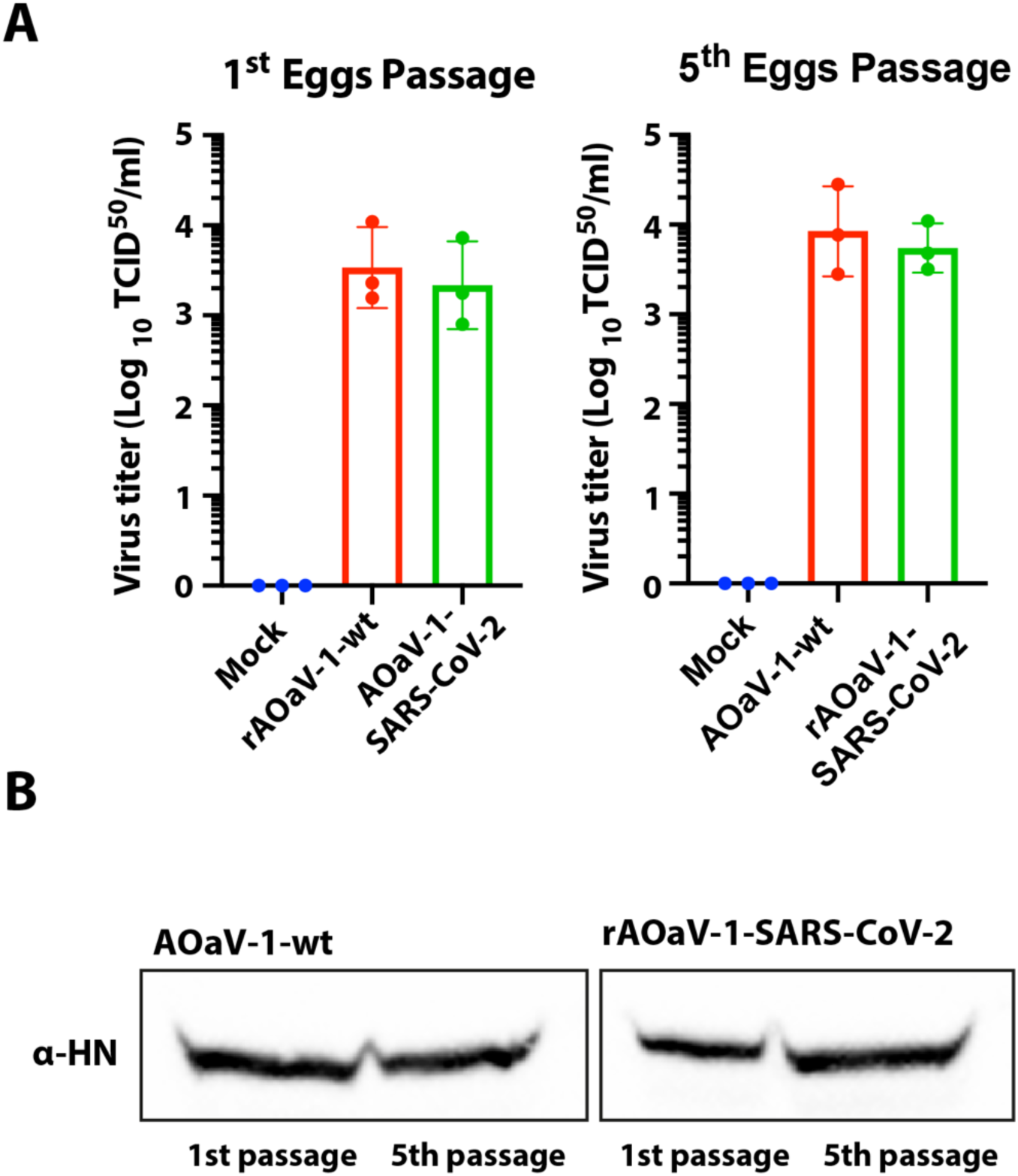
Stability replication of AOaV-1 SARS-CoV-2 vaccine candidate in embryonated chicken eggs. **(A)** The rAOaV-1-SARS-CoV-2 as well as AOaV-1-wt were propagated in eggs and were quantified in Vero cells. The titre quantification, as shown in the bar chart, represent comparable replication. Both rAOaV-1-SARS-CoV-2 and AOaV-1-wt were consecutively propagated in chicken embryonated eggs for 5 passages and the virus titre was quantified in Vero cells. (**B**) The first and fifth passages for both rAOaV-1-SARS-CoV-2 and AOaV-1-wt were used to infect Vero cells for 24 hours before cell lysis and expression analysis for the HN structural protein of the rAOaV-1.

## Discussion

The precipitously increasing deaths, negative impact on lives and livelihood, and trembling economies warrant the urgent development of SARS-CoV-2 vaccine which would be safe, efficacious and scalable. Amongst experimental vaccines being presented for SARS-CoV-2, the vectored-based vaccines hold potential for effective vaccine against CoVID-19^3^. However, each viral vector inherit multiple advantages and disadvantages and careful consideration of gene delivery system may pave the way for an effective vaccine.

We propose here a pre-tested vaccine vector based on the recombinant apathogenic strain of AOaV-1 (i.e. NDV). We engineered AOaV-1 to express the full-length S glycoprotein of SARS-CoV-2 which is a vital viral neutralization and major protective antigen of the virus^24^. Using exhaustive *in vitro* characterization of recombinant vaccine candidate, rAOaV-1-SARS-CoV-2 demonstrated efficient co-expression of surface proteins of AOaV-1 and SARS-CoV-2 in mammalian cell line. This host-range restricted replication in mammals is one of the most attractive properties of AOaV-1^25^. While the mechanism of host range-restriction needs investigations, it is known that AOaV-1 induces a strong interferon response in mammalian cells which in turn limit its replication^26,27^. Additionally, sialic acid receptors for the viral attachment might show fundamental differences between avian and mammalian hosts and thus define the host restriction. It has been observed that bovine parainfluenza virus type 3, another paramyxovirus in the same family, determine the host range restriction through multiple viral proteins^28^. Notwithstanding, AOaV-1 are clearly highly restricted to primates and this selective replication is irrespective of any of the known pathotypes such as mesogenic or lentogenic strains of AOaV-1^29^.

In addition to above mentioned factors, the AOaV-1 carries a range of advantages over multiple other vectors. The AOaV-1 are antigenically distinct from viruses that are known to naturally infect human and AOaV-1 don’t show cross-reactivities with antibodies raised against other paramyxoviruses. Therefore, AOaV-1 induces strong immune responses in the human population. Pertinent to this feature and interferon sensitivity, AOaV-1 are considered a valuable oncolytic agent and thus propose safety in human upon parenterally administration^30^. www.cancer.gov/cancertopics/pdq/cam/NDV/healthprofessional. Potential seroconversion and a transient conjunctivitis may occur in poultry health workers, which subsides within four days without any additional or systemic symptoms^31^. Altogether, AOaV-1 presents a pre-tested and proven safety profile in human and may highlight speeding rolling out of the recombinant vaccines in general human population.

Similar to other paramyxoviruses, AOaV-1 enters host cells by direct fusion at the plasma membrane through a pH-independent mechanism^32,33^. The AOaV-1 can also enter host cells by an endocytic pathway. The entry of the AOaV-1 in the cell is then mediated by the surface fusion (F) glycoprotein, which is also a major determinant of AOaV-1 virulence in birds^34^. The viral infectivity requires cleavage of the F protein through the intracellular ubiquitous proteases including furin, allowing disseminated replication in multiples organs and tissues. A mechanism similar to AOaV-1, rAOaV-1-SARS-CoV-2 entry the cells through proteolytic cleavage of the S protein^24^. The trypsin-independent infectivity of rAOaV-1-SARS-CoV-2, as was observed in our study, facilitated the vector propagation in cells culture model as well as embryonated chicken eggs.

It has previously been shown that insertion of the transgene may allow the vector sensitive to neutralization by antibodies which are specific to the inserted protein and may facilitate the seepage of neutralization by vector-specific neutralizing antibodies^25,35^. The rAOaV-1-SARS-CoV-2 expresses both the native structural HN protein as well as S glycoprotein and therefore the entry into the cell may be attributed to the AOaV-1-like or SARS-CoV-2-like. In conjunction to this hypothesis, the anti-AOaV-1 antiserum as well as anti-SARS-CoV-2 could substantially block the entry of the rAOaV-1-SARS-CoV-2. Additionally, the recombinant AOaV-1-SARS-CoV-2 offers an exciting system to underpin functional interactions between native and foreign envelope glycoproteins in one viral particle.

Recently, insertion of the foreign genes in the backbone of the AOaV-1 has been practiced for multiple viruses including influenza, SARS, human immunodeficiency virus^7,35^, human parainfluenza, rabies^9^, Nipah disease^10^, Rift Valley fever^11^, Ebola and highly pathogenic H5N1^12,13^. In several studies, it has been demonstrated that insertion of the transgene into the genomes of AOaV-1 resulted in reduced pathogenicity in poultry birds. This safety was maintained even after the expression of the HA gene from a highly pathogenic avian influenza virus^11-13^. Sustained, stable and progressive replication of both wt and recombinant viruses demonstrated effective spread within cells independently.

Taken together, our results demonstrated that the recombinant vector expressing S protein propagated stably in chicken embryonated eggs for several consecutive passages, and this expression did not significantly impact the *in vitro* growth characteristics of the recombinants. The presented respiratory vaccine candidate has the potential for further development as vaccine vector to be available for expedited vaccine development for pre-clinical and clinical studies.

## Material and Methods

### Cells and viruses

AOaV-1-wt vaccine vector was rescued as described previously^36^, passaged twice on African green monkey kidney (VERO) cells, and in embryonated chicken eggs. Vero cell line was obtained from the American Type Culture Collection (ATCC, Manassas, VA). These cells were grown in Dulbecco’s minimal essential medium (DMEM) containing 10% fetal bovine serum (FBS). Fertile embryonated chicken eggs were purchased from Henry Stewart & Co Ltd, UK.

### Generation of recombinant AOaV-1 expressing S protein of SARS-CoV-2

The genome of AOaV-1 encodes six structural genes in the order of nucleocapsid (N), phosphoprotein (P), matrix (M), fusion (F), hemagglutinin-neuraminidase (HN) and RNA dependent RNA polymerase (RdRP, also known as L). The transcription of the linear gene starts from gene start (GS) and ends at the gene-end (GE). Between the GS and GE, there is an intergenic sequence (IG). The foreign gene can be inserted at any of the gene junction with variable level of transcription; however, recent studies have assessed the optimal gene expression when inserted between P and M gene^35,37,38^. This arrangement of the gene is identical among all AOaV-1.

In order to offer a competitive reverse genetic system as a novel vaccine vector, an avirulent strain of AOaV-1 was used carrying lentogenic-alike (e.g. NDV) cleavage site and pathogenicity (Patent pending on the technology). The full length antigenomic sequence of AOaV-1, originally isolated from asymptomatic wild birds, was partially *de novo* synthesised (NBS Biologicals, UK) using a novel sequence modification approach. Rest of the genome length mainly constituting around the P gene was cloned using overlapping PCRs. The entire cassette was then shuttled into the TVT7R(0,0) (Addgene Plasmid #98631). In order to avoid incorporation of an extra non-viral gene residues into the transcribed minigenome and efficient transcription of the gene, an autocatalytic and “rule of six” adhered HHRz was introduced at the 5’-end and hepatitis delta virus ribozyme (HdvRz) at the 3’ end of the antigenome.

An expression cassette for the full-length S gene was first *in silico* generated, containing the Kozak sequence, GE, GS, IG and the ORF for the full length S gene was codon-optimized for *homo sapiens* codon usage and inserted into the unique *PmeI* site between P and M genes, which was originally preserved during the cloning of complete antigenome. The construct was named rAOaV-1-SARS-CoV-2 whereas the AOaV-1 without the insertion of the foreign gene (AOaV-1-wt) was used as infection control throughout the study. The orientation of the inserted S gene was confirmed by the nucleotide sequence analysis at the time of cloning as well as during the propagation of the vector in cells and chicken embryonated eggs.

The rAOaV-1-SARS-CoV-2 and AOaV-1-wt were used to rescue the infectious viruses as described previously^36^ with substantial modifications. Briefly, Vero cells were infected with Modified vaccinia Ankara (MVA) expressing the T7 polymerase at a multiplicity of infection 1.0 for 6 hours. These cells were transfected with Lipofectamine 2000 using rAOaV-1-SARS-CoV-2 and AOaV-1-wt backbones as well as supporting N, P and L genes expression plasmids (ratio of 1:0.8:0.4:0.1) for 72 hours. After 3 days post-infection, cells and cell supernatants were mixed and freeze-thaw for three time at -80 before inoculating into 8-days-old embryonated chicken eggs. After additional three days, individual eggs were screened using hemagglutination assay and real-time PCR as we described before^39-41^. Successfully rescued isolates were further propagated to generate viral stock and for *in vitro* characterization.

### Propagation of viruses in eggs and cells

The infectivity of the recombinant virus and parental wild type strains were characterized using a standard hemagglutination assay (HA). The 50% tissue infectious dose (TCID_50_) assay on Vero cells in 96-well plates, was performed and calculated following standard procedures^39,40,42^.

### Western blotting

To confirm expression of the viral protein, Vero cells were infected with recombinant and wild type viruses at a multiplicity of infection (moi) of 0.1 as we described earlier^43^. Cell lysates were collected at 24 h post infection, subjected to Western blot. Briefly, after 24h post-infection, cells were washed one time with phosphate buffered saline (PBS) followed by adding 100 μL ice-cold NP40 lysis buffer (completed with protease inhibitors cocktail) per well and incubated for 30 min on ice. All cell lysates were denatured at 98°C for 8 min, proteins were separated by sodium dodecyl sulphate polyacrylamide gel electrophoresis (SDS-PAGE), and subsequently transferred to nitrocellulose membranes^43^. The cell lysates were subjected for centrifugation at 13,000 rpm for 30 min at 4°C and the supernatant were incubated in sample loading buffer 2x (Biorad) containing 10% β-mercaptoethanol for 5 min at 98 °C, and separated on a 10% sodium dodecyl sulfate–polyacrylamide gel electrophoresis (SDS-PAGE), and proteins were transferred onto polyvinylidene difluoride (PVDF) membranes. The S and HN viral proteins were detected by incubation with primary antibodies (1:500 dilution). After probing with primary antibodies, the blots were incubated with peroxidase-conjugated species-specific secondary antibodies (Abcam) and visualised by chemiluminescence (Chemidoc, BioRad, Hercules, CA, USA), as we performed earlier^43^.

### Immunofluorescence

The expression of the S and HN proteins in recombinant virus-infected cells were evaluated by immunofluorescence assays using confocal microscopy. Vero cells grown on coverslips in 24-well plates, were infected with wild type or recombinant viruses or viruses pre-incubated with antisera for 24h. After fixing the cells with 4% paraformaldehyde in PBS for 30 min, washed with PBS, and permeabilized with 0.1% Triton X-100 for 10 min. After blocking of the cells with 5% bovine serum albumin (BSA) in PBS, they were incubated with monoclonal antibody (mAb)^44^ to probe HN or S or both proteins. Binding of primary antibodies were visualized using Alexa 488 α-rabbit and 568 α-mouse secondary antibodies (Invitrogen). The S and HN proteins expression were analysed through fluorescence for wild-type and recombinant viruses compared to mock infected cells. The 40,6-diamidino-2-phenylindole (DAPI) was used to stain cell nuclei and the images were captured using a Zeiss confocal laser-scanning microscope (Zeiss, Kohen, Germany). Digital images were processed using Adobe illustrator software, and the same parameters were applied to the whole image area.

### Stability and growth characteristics of NDV constructs using real-time PCR

The stability of recombinant vaccine candidate compared to parental wild type were grown sequentially in embryonated chicken eggs for at least 5 passages. RNA was extracted from the allantoic fluid of recombinant and wild type after the first and fifth passages in eggs using the QIAamp^®^ Viral RNA Mini Kit (Qiagen). The real-time qRT-PCR was performed using *SuperScript*™ *III Platinum*™ *One*-*Step qRT*-*PCR Kit* to detect NDV M gene^41^, which enabled the calculation of viral genome copies that was plotted against hours post-infection for each of the viruses to produce standard curves. To investigate the growth properties of the recombinant viruses, Vero cells were infected in 6-well plates at a moi of 0.1 after washed twice with PBS and an adsorption time of 60 min. Cell supernatants were harvested 0, 6, 12, 24 and 48h after infection (p.i). Viral titers (50% tissue culture infectious dose, TCID_50_) were calculated by IFA using primary antibodies against viral proteins and Alexa FluorTM 488 α-rabbit and Alexa FluorTM 586 α-mouse as secondary antibodies (Invitrogen), respectively.

### Statistical analysis

Virus-infected and control groups were compared using Student’s *t*-test. All statistical analyses and figures were conducted in the GraphPad Prism (GraphPad Software, La Jolla, CA, USA).

## Acknowledgements

This study was supported by the Biotechnology and Biological Sciences Research Council (BBSRC) grant BB/M008681/2.

## Authors contribution

M.M. designed the experiment; M.R. and M.M. collected data; M.R. and M.M. performed data analysis and wrote the manuscript; M.R. and M.M. discussed results, revised the manuscript substantively. M.M. reviewed and approved the manuscript.

## Competing interests

The authors declare no competing interests.

## Supplementary Information

Supplementary figures are associated with this manuscript.

## Supplementary Figure Legends

**Supplementary Figure 1**. A real-time PCR-based detection of rescued AOaV-1-wt (**A**) and rAOaV-1-SARS-CoV-2 (**B**) in chicken embryonated eggs. (**C**) Quantitative presentation of AOaV-1-wt or rAOaV-1-SARS-CoV-2 detection. (**D**) Hemagglutination assay used to screen the chicken embryonated eggs for the presence (+)/absence (-) of AOaV-1-wt or rAOaV-1-SARS-CoV-2.

**Supplementary Figure 2. Replication of wt and recombinant viruses in Vero cells. (A)** Expanded confocal microscopic images representing Figure 2 A-C in the main text. (**B**) Comprehensive and expanded confocal microscopic images representing Figure 2 D-F in the main text.

**Supplementary Figure 3:** Raw Western blot images used in Figure 1, 4 and 5. The portion of the figures used in the main manuscript is marked with the black lined square.

## Notes

### Competing Interest Statement

The authors have declared no competing interest.

## References

1. Wu, Z. & McGoogan, J. M. Characteristics of and Important Lessons From the Coronavirus Disease 2019 (COVID-19). Outbreak in China: Summary of a Report of 72 314 Cases From the Chinese Center for Disease Control and Prevention. JAMA. 10.1001/jama.2020.2648 (2020).

2. Coronaviridae Study Group of the International Committee on Taxonomy of Viruses. The species severe acute respiratory syndrome-related coronavirus: Classifying 2019-nCoV and naming it SARS-CoV-2. Nat. Microbiol. 5, 536–544 (2020).

3. WHO. Draft landscape of COVID-19 candidate vaccines. https://www.who.int/who-documents-detail/draft-landscape-of-covid-19-candidate-vaccines (2020).

4. Gao, Q. et al. Rapid development of an inactivated vaccine candidate for SARS-CoV-2. Science. pii: eabc1932. 10.1126/science.abc1932. (2020).

5. Doremalen, N. et al. ChAdOx1 nCoV-19 vaccination prevents SARS-CoV-2 pneumonia in rhesus macaques. https://doi.org/10.1101/2020.05.13.093195 (2020)

6. Kim, S.H. & Samal, S. K. Newcastle Disease Virus as a Vaccine Vector for Development of Human and Veterinary Vaccines. Viruses. 8(7), pii: E183. 10.3390/v8070183 (2016).

7. Khattar, S. K. et al. Newcastle disease virus expressing human immunodeficiency virus type 1 envelope glycoprotein induces strong mucosal and serum antibody responses in Guinea pigs. J. Virol. 85, 10529–10541 (2011).

8. Carnero, E. et al. Optimization of human immunodeficiency virus gag expression by newcastle disease virus vectors for the induction of potent immune responses. J. Virol. 83, 584–597 (2009).

9. Ge, J. et al. Newcastle disease virus-vectored rabies vaccine is safe, highly immunogenic, and provides long-lasting protection in dogs and cats. J. Virol. 85, 8241–8252 (2011).

10. Kong, D. et al. Newcastle disease virus-vectored Nipah encephalitis vaccines induce B and T cell responses in mice and long-lasting neutralizing antibodies in pigs. Virology 432, 327–335 (2012).

11. Kortekaas, J. et al. Rift Valley fever virus immunity provided by a paramyxovirus vaccine vector. Vaccine. 28, 4394–4401 (2010a).

12. DiNapoli, J. M. et al. Newcastle disease virus-vectored vaccines expressing the hemagglutinin or neuraminidase protein of H5N1 highly pathogenic avian influenza virus protect against virus challenge in monkeys. J. Virol. 84, 1489–1503 (2010a).

13. Ge, J. et al. Newcastle disease virus-based live attenuated vaccine completely protects chickens and mice from lethal challenge of homologous and heterologous H5N1 avian influenza viruses. J. Virol. 81, 150–158 (2007).

14. Khattar SK, Collins PL, Samal SK: Immunization of cattle with recombinant Newcastle disease virus expressing bovine herpesvirus-1 (BHV-1) glycoprotein D induces mucosal and serum antibody responses and provides partial protection against BHV-1. Vaccine 2010, 28:3159–3170.

15. Kortekaas, J. et al. Intramuscular inoculation of calves with an experimental Newcastle disease virus-based vector vaccine elicits neutralizing antibodies against Rift Valley fever virus. Vaccine. 28, 2271–2276 (2010b)

16. DiNapoli, J. M., Yang, L., Samal, S. K., Murphy, B. R., Collins, P. L. & Bukreyev, A. Respiratory tract immunization of non-human primates with a Newcastle disease virus-vectored vaccine candidate against Ebola virus elicits a neutralizing antibody response. Vaccine. 29, 17–25 (2010b)

17. Bukreyev, A. et al. Recombinant newcastle disease virus expressing a foreign viral antigen is attenuated and highly immunogenic in primates. J. Virol. 79, 13275–13284 (2005).

18. Bukreyev, A. & Collins, P. L. Newcastle disease virus as a vaccine vector for humans. Curr. Opin. Mol. Ther. 10, 46–55 (2008).

19. Pecora, A. L. et al. Phase I trial of intravenous administration of PV701, an oncolytic virus, in patients with advanced solid cancers. J. Clin. Oncol. 20, 2251–2266 (2002).

20. Ockert, D. et al. Newcastle disease virus infected intact autologous tumor cell vaccine for adjuvant active specific immunotherapy of resected colorectal carcinoma. Clin. Cancer Res. 2, 21–28 (1996).

21. Karcher, J. et al. Antitumor vaccination in patients with head and neck squamous cell carcinomas with autologous virus-modified tumor cells. Cancer Res. 64, 8057–8061 (2004).

22. Peeters, B. P., de Leeuw, O. S., Koch, G. & Gielkens, A. L. Rescue of Newcastle disease virus from cloned cDNA: evidence that cleavability of the fusion protein is a major determinant for virulence. J. Virol. 73, 5001–5009 (1999).

23. Panda, A., Huang, Z., Elankumaran, S., Rockemann, D. D. & Samal, S. K. Role of fusion protein cleavage site in the virulence of Newcastle disease virus. Microb. Pathog. 36, 1–10 (2004).

24. Hoffmann, M., Kleine-Weber, H. & Pöhlmann, S. A. Multibasic Cleavage Site in the Spike Protein of SARS-CoV-2 Is Essential for Infection of Human Lung Cells. Mol. Cell. pii: S1097-2765(20)30264-1. 10.1016/j.molcel.2020.04.022 (2020).

25. DiNapoli, J. M. et al. Newcastle disease virus, a host range-restricted virus, as a vaccine vector for intranasal immunization against emerging pathogens. Proc. Natl. Acad. Sci. USA. 104(23), 9788–93 (2007).

26. Blach-Olszewska, Z. Interferon induction by Newcastle disease virus in mice. Arch. Immunol. Ther. Exp. (Warsz) 18, 418–441. 62 (1970)

27. Brehm, G. & Kirchner, H. Analysis of the interferons induced in mice in vivo and in macrophages in vitro by Newcastle disease virus and by polyinosinic-polycytidylic acid. J. Interferon Res. 6, 21–28 (1986)

28. Skiadopoulos, M. H. et al. Determinants of the host range restriction of replication of bovine parainfluenza virus type 3 in rhesus monkeys are polygenic. J. Virol. 77, 1141–1148 (2003)

29. Bukreyev, A. et al. Recombinant newcastle disease virus expressing a foreign viral antigen is attenuated and highly immunogenic in primates. J. Virol. 79, 13275–13284 (2005)

30. Lu, L. et al. Immunological characterization of the spike protein of the severe acute respiratory syndrome coronavirus. J. Clin. Microbiol. 42, 1570–1576 (2004)

31. Nelson, C. B., Pomeroy, B. S., Schrall, K., Park, W. E. & Lindeman, R. J. Am J Public Health. 42, 672–678 (1952)

32. Hernandez LD, Hoffman LR, Wolfsberg TG, White JM: Virus-cell and cell-cell fusion. Annu. Rev. Cell Dev. Biol. 12, 627–661 (1996)

33. Lamb, R. A. Paramyxovirus fusion: a hypothesis for changes. Virology. 197, 1–11 (1993).

34. Cantin, C., Holguera, J., Ferreira, L., Villar, E. & Munoz-Barroso, I. Newcastle disease virus may enter cells by caveolae-mediated endocytosis. J. Gen. Virol. 88, 559–569 (2007)

35. Carnero, E., Li, W., Borderia, A. V., Moltedo, B., Moran, T. & Garcia-Sastre, A. Optimization of human immunodeficiency virus gag expression by newcastle disease virus vectors for the induction of potent immune responses. J. Virol. 83, 584–597 (2009)

36. Ayllon, J., García-Sastre, A. & Martínez-Sobrido, L. Rescue of recombinant Newcastle disease virus from cDNA. J. Vis. Exp. (80) 10.3791/50830. (2013).

37. Zhao, W., Zhang, Z., Zsak, L. & Yu, Q. P and M gene junction is the optimal insertion site in Newcastle disease virus vaccine vector for foreign gene expression. J. Gen. Virol. 96(1), 40–45 (2015).

38. Qingzhong Yu, Q. Have we found an Optimal Insertion Site in a Newcastle Disease Virus Vector to Express a Foreign Gene for Vaccine and Gene Therapy Purposes? Br. J. Virol. 2(2), 15–18 (2015)

39. OIE. Newcastle Disease, Biological Standards Commission, Manual of Diagnostic Tests and Vaccines for Terrestrial Animals: Mammals, Birds and Bees. (7th ed), World Organisation for Animal Health, Paris, France, pp. 555–574 (2012).

40. Grimes, S. E. A Basic Laboratory Manual for the Small-Scale Production and Testing of 1-2 Newcastle Disease Vaccine. FAO Regional Office for Asia and the Pacific (RAP); Bangkok, Thailand (2002).

41. Wise, M. G. et al. Development of a real-time reverse-transcription PCR for detection of Newcastle disease virus RNA in clinical samples. J. of Clin. Microbiol. 42, 329–338 (2004).

42. Reed, L. & Muench, H. A simple method of estimating fifty per cent endpoints. Am. J. Epidemiol. 27, 493–497 (1938).

43. Atasoy, M. O, Rohaim, M. A & Munir, M. Simultaneous Deletion of Virulence Factors and Insertion of Antigens Into the Infectious Laryngotracheitis Virus Using NHEJ-CRISPR/Cas9 and Cre-Lox System for Construction of a Stable Vaccine Vector. Vaccines, 5, 7, 207 (2019)

44. Russell, P. H. et al. The Characterization of Monoclonal Antibodies to Newcastle Disease Virus. J. Gen. Virol. 64, 2069–2072 (1983).

